# Loss of *Hnf1b* in differentiated proximal tubule cells uncovers nephron segment plasticity

**DOI:** 10.64898/2026.08.12.744527

**Authors:** Zeinab Dehghani-Ghobadi, Eunah Chung, Azadeh Haghighitalab, Mohammed Sayed, Christopher Ahn, Yueh-Chiang Hu, Hee-Woong Lim, Joo-Seop Park

## Abstract

HNF1B is a transcription factor required for proximal tubule (PT) specification during kidney development, but whether it is also required to maintain PT identity after differentiation remains unknown. Using PT-specific genetic deletion in mice, we found that loss of *Hnf1b* in differentiated PT cells causes cyst formation and early postnatal lethality. PT-specific transcriptomic analysis revealed downregulation of PT-specific gene programs, including *Hnf4a* and PT-enriched transport and metabolic genes. Strikingly, *Hnf1b*-deficient PT cells ectopically activated podocyte-specific genes, including *Wt1* and *Nphs1*, demonstrating that PT cells retain the capacity to engage alternative nephron segment programs when identity-stabilizing mechanisms are disrupted. In addition, loss of *Hnf1b* disrupted epithelial integrity, as evidenced by reduced epithelial adhesion gene expression and induction of mesenchymal markers. Wnt/β-catenin signaling was also aberrantly activated, suggesting broader dysregulation of epithelial homeostasis. These findings establish HNF1B as a critical post-specification regulator of PT identity that sustains PT-specific transcriptional programs and actively suppresses alternative segmental identity programs.

## INTRODUCTION

During mammalian kidney development, mesenchymal nephron progenitors (mNPs) become epithelialized to form renal vesicles, which subsequently differentiate into the major nephron segments, including podocytes, parietal epithelial cells, proximal tubule (PT), loop of Henle, distal tubule, and connecting tubule (1, 2). The PT, the largest nephron segment, performs the majority of solute and water reabsorption, a process essential for systemic homeostasis (3–5). To sustain these functions, PT cells maintain high metabolic activity and a transcriptional program enriched for transporters and metabolic enzymes (3, 6–8). Loss of PT differentiation is observed in kidney disease and maladaptive repair, highlighting the importance of understanding the mechanisms that actively sustain PT identity (9–11). While the mechanisms that specify nephron segments during development have been studied (2, 12), the transcriptional programs that actively maintain PT identity remain poorly understood.

HNF1B is a key transcription factor required for nephron segmentation (13–17). Heterozygous mutations in *HNF1B* are among the most common monogenic causes of congenital kidney disease in humans, leading to renal cysts, hypoplasia, and tubulointerstitial disease, often referred to as maturity-onset diabetes of the young type 5 (MODY5) or renal cysts and diabetes (RCAD) syndrome (18–20). In both mouse and zebrafish, *Hnf1b* is required for proper proximo-distal patterning of the nephron during development (13, 14, 16, 17). Mouse studies defining this role have relied on early deletion models driven by *Six2Cre* or *Wnt4Cre*, which delete *Hnf1b* before or during epithelialization of mNPs (14, 16, 17). These models demonstrate that loss of *Hnf1b* causes glomerulocyst formation, disrupts development of mature nephron segments, and blocks expression of HNF4A, a transcription factor required for PT differentiation (6, 21). In addition, studies using *KspCre* or *Pkhd1Cre* have established important roles for *Hnf1b* in the collecting duct epithelium (22–33). Collectively, these studies have largely focused on *Hnf1b* function either during early nephrogenesis or in the collecting duct, leaving its role in maintaining segment-specific gene expression programs in differentiated PT cells largely unexplored.

To address this gap, we generated a mouse model in which *Hnf1b* is conditionally deleted from PT cells after their formation using the PT-specific *Slc34a1Cre* driver (34). Unlike previously used Cre drivers, *Slc34a1Cre* targets PT cells after segmental identity has been established, enabling direct assessment of *Hnf1b* function in differentiated PT cells. We isolated PT cells by fluorescence-activated cell sorting (FACS) and performed transcriptomic profiling, histological characterization, and lineage tracing. This strategy enables direct assessment of whether *Hnf1b* is required to sustain PT identity after segment specification. Here, we show that loss of *Hnf1b* in differentiated PT cells leads to pronounced disruption of PT-specific transcriptional programs and ectopic activation of podocyte-associated genes, revealing an unexpected degree of nephron segment plasticity. These findings demonstrate that PT identity is not a fixed endpoint of differentiation but instead requires active transcriptional maintenance and suppression of alternative nephron segment programs.

## RESULTS

### HNF1B maintains PT identity and suppresses ectopic podocyte gene programs

In the developing mouse kidney, *Hnf1b* is expressed in all renal epithelial cells except podocytes (Supplemental Figure 1A and 1B). Although *Hnf1b* is required for *Hnf4a* expression during PT specification (14, 16), it remains unknown whether *Hnf1b* is also required to maintain PT identity after segmental identity has been established. To address this, we conditionally deleted *Hnf1b* using *Slc34a1Cre*, which is expressed specifically in differentiated PT cells in the developing mouse kidney (34). This approach permits *Hnf1b*- dependent activation of *Hnf4a* during PT specification to proceed normally, with deletion occurring only after PT segmental identity is established, thereby isolating the role of *Hnf1b* in identity maintenance from its earlier role in PT specification. Most *Hnf1b* mutant mice developed severe polycystic kidneys and died by postnatal day 10 (Figure 1A). To determine whether cyst-lining cells originated from *Hnf1b*-deficient PT cells rather than other nephron segments, we performed lineage tracing using *Slc34a1Cre*-mediated activation of the *Rosa26- Sun1* GFP reporter. The majority of cyst-lining epithelial cells were Sun1-positive and HNF1B-negative, confirming that the cysts arose from recombined PT cells (Figure 1B).

**Figure 1.**
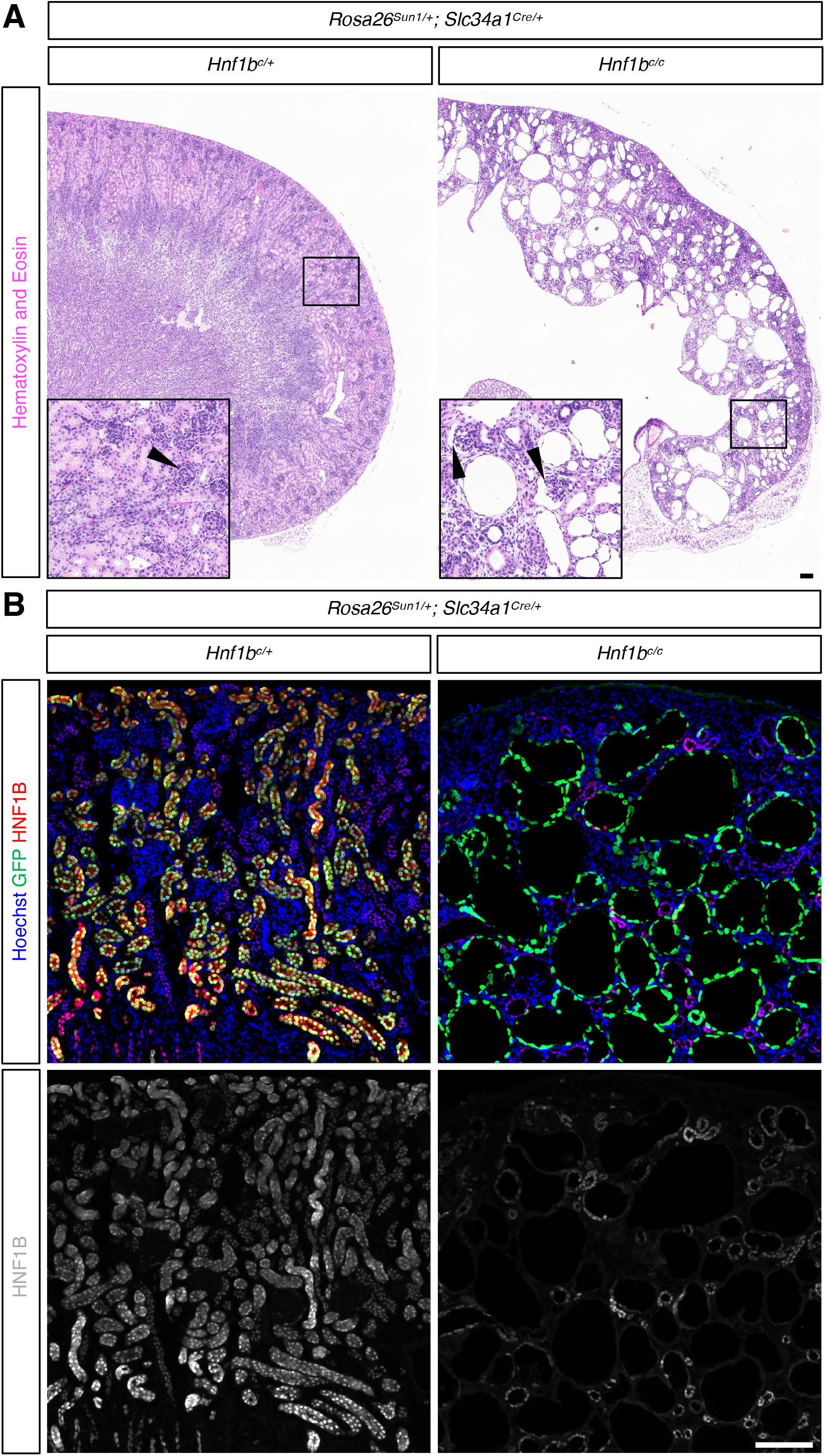
Loss of *Hnf1b* in PT cells leads to a polycystic kidney phenotype. (A) H&E staining shows extensive cyst formation in *Hnf1b* mutant kidneys. Higher-magnification views (insets) of the regions outlined in black reveal normal renal corpuscle morphology in control kidneys, whereas *Hnf1b* mutant kidneys exhibit dilated Bowman’s space (black arrowheads). (B) Lineage tracing with *Slc34a1Cre*-mediated activation of the *Rosa26-Sun1* reporter shows that the cyst-lining epithelial cells originate from *Slc34a1*+ PT cells. These SUN1+ cells lack HNF1B, indicating that cyst formation occurs within *Hnf1b*-deficient PT cells. (A, B) Representative images from four biological replicates per genotype are shown. Stage: postnatal day 9; Scale bar: 100 μm.

To define the transcriptional consequences of *Hnf1b* loss, we performed bulk RNA-seq on FACS-isolated PT cells from *Hnf1b* mutant and littermate control kidneys (Supplemental Table 1). Differential expression analysis identified 4,029 significantly altered genes, comprising 2,212 upregulated and 1,817 downregulated transcripts (fold change >2, FDR <0.01; Supplemental Figure 1C). To determine which nephron cell types most closely matched the transcriptional changes observed in *Hnf1b*-deficient PT cells, we projected module scores derived from differentially expressed genes onto a single-cell RNA sequencing (scRNA-seq) reference dataset of the developing mouse kidney (34–36). Downregulated genes mapped predominantly to the PT cluster, indicating loss of PT-specific transcriptional programs (Supplemental Figure 1D), whereas upregulated genes mapped primarily to the podocyte cluster, suggesting aberrant activation of a podocyte-associated transcriptional program in *Hnf1b*-deficient PT cells (Supplemental Figure 1E). Figure 2 highlights representative differentially expressed genes. Heatmaps demonstrate that these gene expression changes are consistent across both sexes, while dot plot analysis using scRNA-seq data shows that downregulated genes are enriched in PT cells and their progenitors, whereas upregulated genes are preferentially expressed in podocytes and their progenitors. Gene Ontology and KEGG pathway analyses further supported these findings: downregulated genes were enriched for PT metabolic functions, including fatty acid β-oxidation, mitochondrial transport, oxidative phosphorylation, and peroxisome metabolism, while upregulated genes were enriched for extracellular matrix organization, Wnt signaling, and developmental pathways (Supplemental Figure 2).

**Figure 2.**
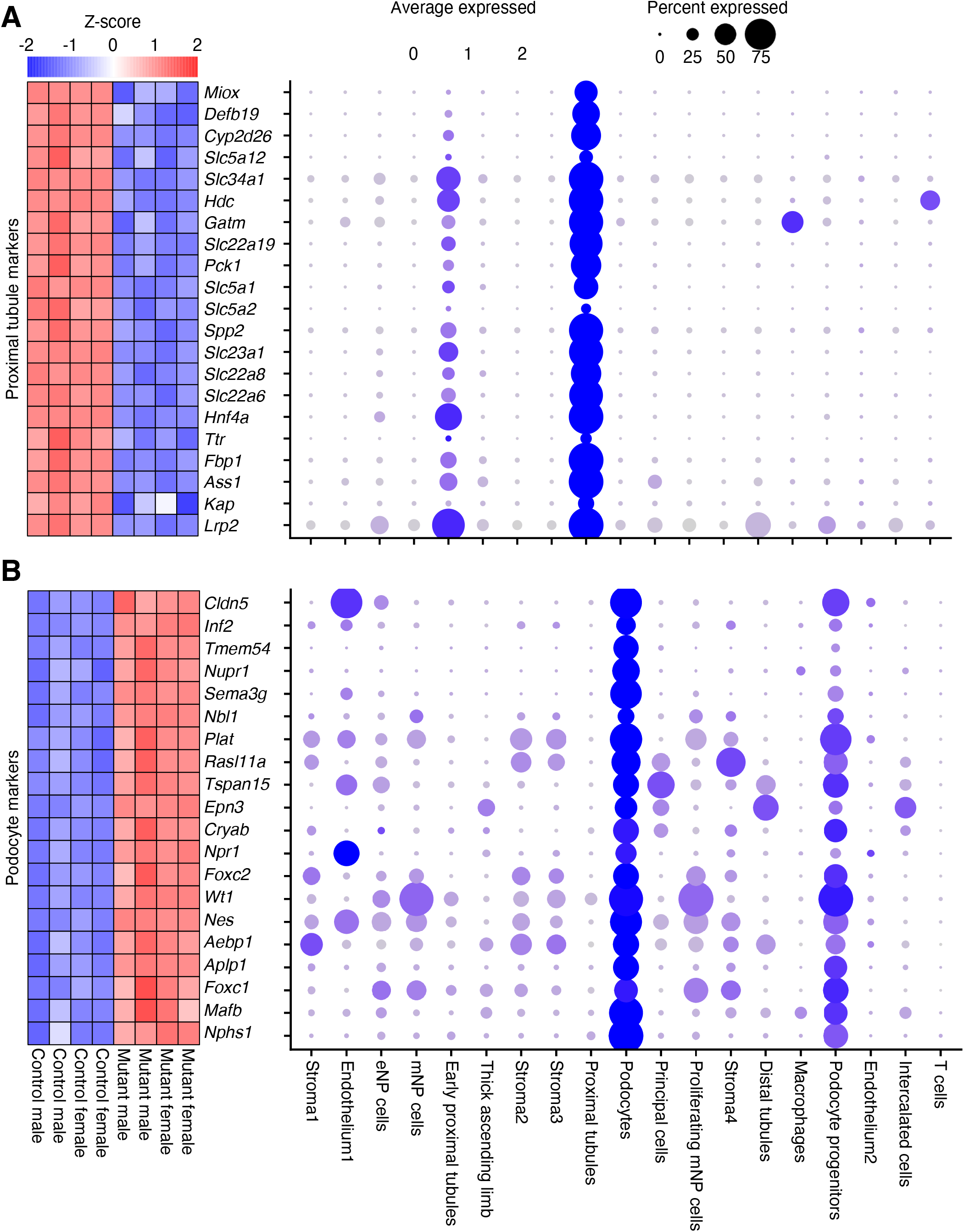
Identification of differentially expressed genes in *Hnf1b*-deficient PT cells. Bulk RNA-seq analysis of FACS-isolated PT cells at postnatal day 1 reveals differentially expressed genes between control and *Hnf1b* mutant kidneys, as shown in the heatmap. In mutant kidneys, PT-enriched genes are downregulated (A), whereas podocyte-associated genes are upregulated (B). These expression changes were observed in both male and female mice. Color intensity represents the magnitude of expression change, with blue indicating decreased and red indicating increased expression. The dot plot displays cell–type–specific expression of marker genes based on previously published single-cell RNA-seq data from mouse kidneys at E18.5 or P0 (GSE214024). Dot color indicates the average expression level of each gene within a given cell type, whereas dot size represents the percentage of cells expressing that gene.

These transcriptional changes were validated at the cellular level. Immunofluorescence confirmed that canonical PT marker expression, including HNF4A, ASS1, LRP2, MIOX, PCK1, GATM, SLC34A1, and SLC5A2, was markedly diminished or absent in Sun1-positive *Hnf1b*-deficient PT cells, along with loss of *Lotus tetragonolobus* lectin (LTL) staining (Figure 3). PT marker expression was retained in a subset of Sun1- negative cells, indicating escape from Cre-mediated recombination and retention of HNF1B expression in a minority of PT cells. Ectopic activation of the podocyte gene program in *Hnf1b*-deficient PT cells was confirmed by immunofluorescence and Hybridization Chain Reaction Fluorescence *In Situ* Hybridization (HCR-FISH). In control kidneys, podocyte marker genes were expressed exclusively in podocytes. In *Hnf1b* mutant kidneys, these markers were aberrantly detected in Sun1-positive PT cells, with ectopic WT1 protein confirmed by immunofluorescence (Figure 4A), ectopic *Nphs1* and *Plat* by HCR-FISH (Figure 4B), and ectopic *Cldn5* and *Mafb* by HCR-FISH (Supplemental Figure 3). Together, these findings identify HNF1B as a critical factor required to maintain PT identity, whose loss disrupts the PT transcriptional program and causes ectopic activation of podocyte-associated gene expression.

**Figure 3.**
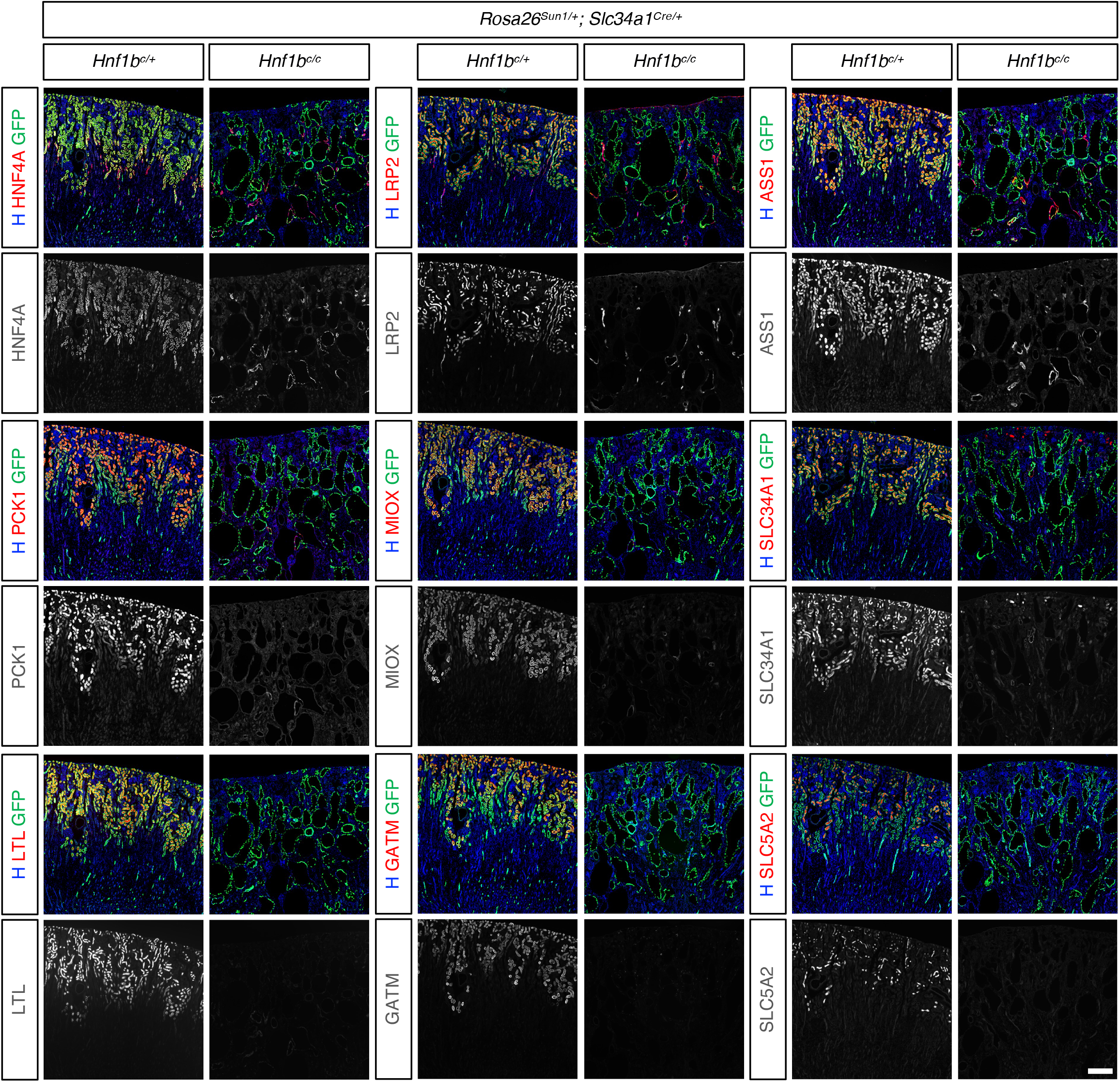
HNF1B is required to maintain PT identity. The mutant kidney exhibits marked reductions in HNF4A expression and PT markers, including ASS1, LRP2, MIOX, PCK1, GATM, SLC34A1, and SLC5A2, together with diminished LTL staining. These findings indicate loss of PT differentiation following *Hnf1b* deletion and are consistent with disruption of HNF4A-dependent transcriptional programs. Representative images from four biological replicates per genotype are shown. H: Hoechst stain; Stage: postnatal day 9; Scale bar: 200 μm.

**Figure 4.**
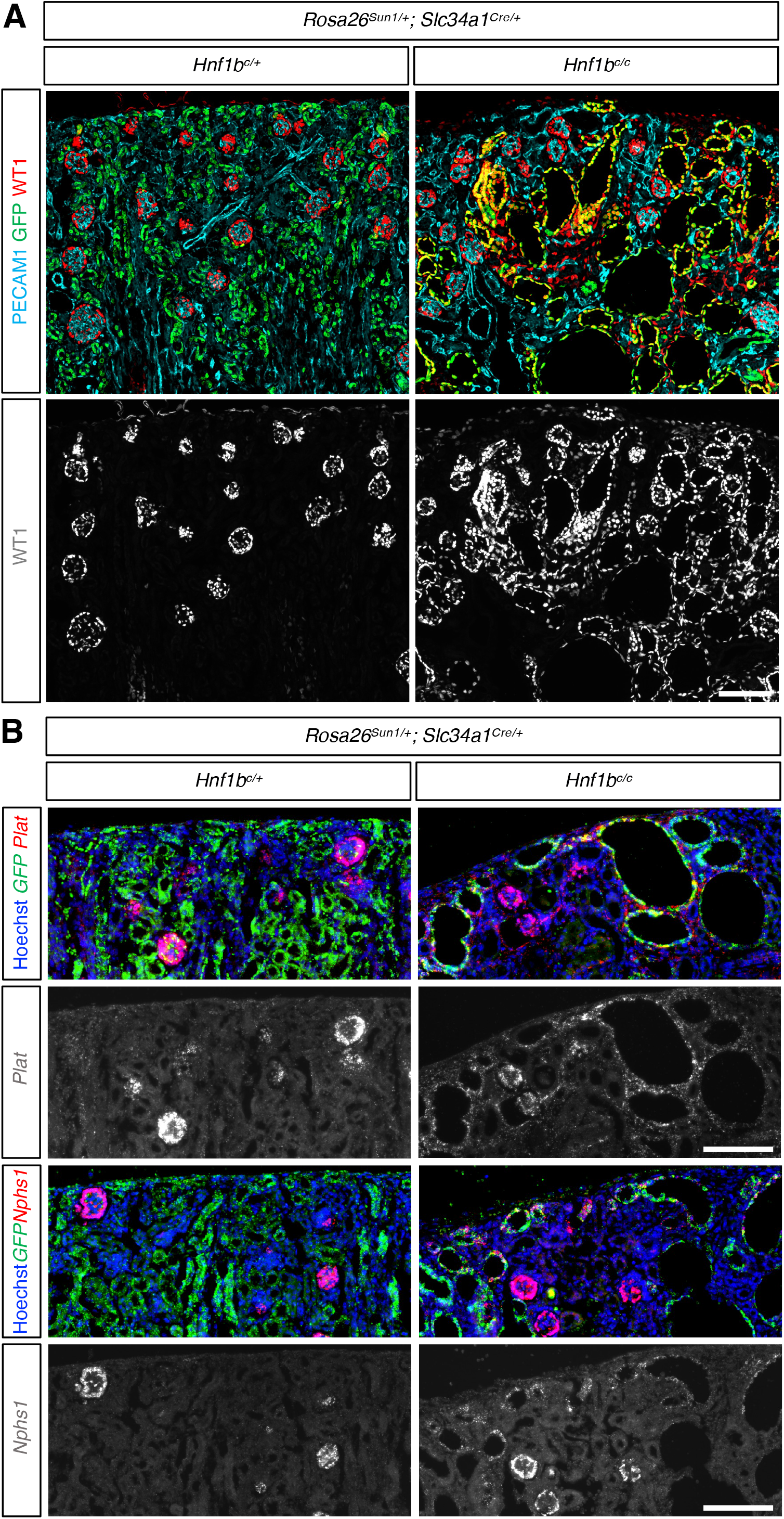
Loss of *Hnf1b* in PT cells induces ectopic activation of podocyte-associated genes. (A) In control kidneys, WT1 expression is restricted to podocytes and absent from most SUN1+ cells. In contrast, *Hnf1b* mutant kidneys exhibit ectopic WT1 expression in SUN1+ PT cells. (B) HCR-FISH demonstrates induction of podocyte-associated genes in *Hnf1b* mutant PT cells. In control kidneys, *Plat* and *Nphs1* expression is restricted to podocytes. In *Hnf1b* mutant kidneys, both genes are ectopically expressed in SUN1+ cells. (A, B) Representative images from four biological replicates per genotype are shown. Stage: postnatal day 9; Scale bar: 100 μm.

### *Slc34a1Cre*-mediated *Hnf1b* deletion causes medullary cysts in PT-derived descending thin limb cells

Following *Hnf1b* deletion, renal cysts were first detected exclusively in the medulla during early postnatal stages (Supplemental Figure 4), with cortical cysts emerging at later time points (Figure 1). This spatiotemporal pattern suggests that cystogenesis initiates within the medulla and subsequently progresses to the cortex, implying that specific medullary cell populations may be particularly susceptible to cyst formation. Consistent with this, prior lineage-tracing studies demonstrated that *Slc34a1Cre* targets not only PT cells but also a subset of descending thin limb (DTL) cells, which are derived from PT cells during kidney development (34). To define the segmental identity of medullary cyst-lining epithelial cells, we analyzed nephron segment-specific marker expression. Cyst-lining cells were negative for SLC12A1, AQP2, and GATA3, excluding thick ascending limb and collecting duct identities. Of the markers examined, only AQP1 was detected. However, AQP1 expression was restricted to a subset of Sun1-positive cyst-lining cells, whereas the majority were AQP1-negative (Supplemental Figure 5A), suggesting that medullary cyst-lining cells do not uniformly retain DTL identity.

Because loss of AQP1 expression could reflect disruption of DTL differentiation rather than exclusion of a DTL origin, we next examined whether *Hnf1b* deletion affects the *Hnf1b-Hnf4a-Aqp1* regulatory axis in PT-derived DTL cells. Our previous work demonstrated that a subset of DTL cells is derived from PT cells and that loss of *Hnf4a* impairs AQP1 expression in the DTL (34). Because *Hnf1b* is required for *Hnf4a* expression, we predicted that *Hnf1b* deletion using *Slc34a1Cre* would similarly disrupt AQP1 expression in PT-derived DTL cells. To test this, we performed co-immunostaining for AQP1, PECAM1, and Sun1. Because AQP1 protein is expressed in both the DTL and vasa recta within the renal papilla (34, 37), this approach enabled discrimination of vascular structures (AQP1-positive, PECAM1-positive) from DTL epithelium, while simultaneously identifying recombined (Sun1-positive) and non-recombined (Sun1-negative) cells. Consistent with this regulatory relationship, a subset of recombined PT-derived DTL cells retained AQP1 expression, whereas the majority lost AQP1 expression following *Hnf1b* deletion (Supplemental Figure 5B), indicating compromised DTL identity in the absence of *Hnf1b*.

### Loss of *Hnf1b* disrupts epithelial polarity and promotes proliferative remodeling

Because epithelial polarity is critical for maintenance of tubular architecture and is frequently disrupted in cystic kidney disease (38, 39), we examined whether these processes were affected following *Hnf1b* deletion in PT cells. Immunostaining demonstrated loss of the adherens junction protein E-cadherin (CDH1) in *Hnf1b*- deficient PT cells, accompanied by increased expression of the mesenchymal markers Vimentin (VIM) and Transgelin (TAGLN) within mutant tubules and adjacent stromal cells (Figure 5). Assessment of polarity markers further revealed disruption of apical-basal organization. In Sun1-positive PT cells in control kidneys, PARD6B localized to the apical domain, whereas LAMA1 formed a continuous basement membrane (Supplemental Figure 6). In contrast, *Hnf1b*-deficient Sun1-positive PTs lacked apical PARD6B and exhibited reduced or mislocalized LAMA1 staining , indicating impaired epithelial polarity and basement membrane organization (Supplemental Figure 6).

**Figure 5.**
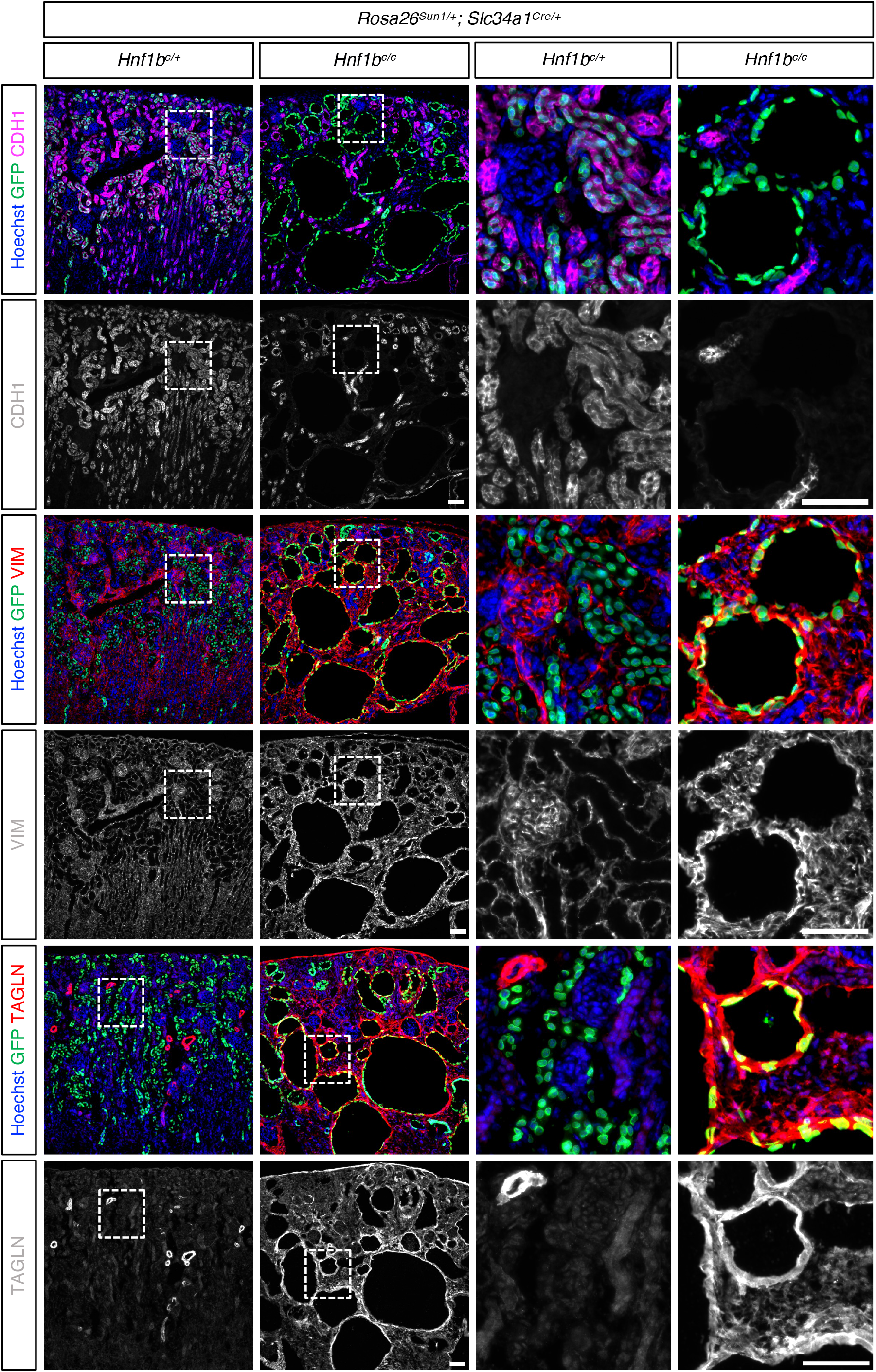
PT-specific deletion of *Hnf1b* reduces CDH1 expression and induces mesenchymal marker expression. Immunostaining of mutant kidneys shows reduced expression of the epithelial marker CDH1 (E- cadherin) in SUN1+ PT cells, together with induction of the mesenchymal markers VIM (Vimentin) and TAGLN (Transgelin) in SUN1+ PT cells and adjacent stromal cells. The regions outlined by dotted boxes in the left panels are shown at higher magnification in the corresponding panels on the right. Representative images from four biological replicates per genotype are shown. Stage: postnatal day 9; Scale bar: 50 μm.

Because epithelial dysfunction in cystic kidney disease is commonly associated with increased proliferation (40), we next examined cell proliferation in mutant kidneys. Ki67 staining demonstrated increased proliferation following *Hnf1b* deletion, including within Sun1-positive PT-derived epithelial cells and cortical nephrogenic regions, and quantification confirmed a significant increase in the percentage of Sun1-positive cells co-expressing Ki67 compared with controls (Supplemental Figure 7A). RNA-seq analysis additionally identified increased expression of *Igf2*, a mitogenic growth factor, in *Hnf1b*-deficient PT cells (Supplemental Table 1). Consistent with a potential role in driving epithelial proliferation, HCR-FISH revealed minimal *Igf2* expression in control kidneys, whereas mutant kidneys exhibited increased *Igf2* signal in Sun1-positive PT-derived cells and cortical nephrogenic regions, with quantification confirming a significant increase in Igf2-positive area compared with controls (Supplemental Figure 7B). Together, these findings demonstrate that loss of *Hnf1b* disrupts epithelial organization, drives a proliferative epithelial response, and induces expression of the mitogenic factor *Igf2* in PT-derived cells.

### Loss of *Hnf1b* in PTs activates Wnt-associated transcriptional programs and stromal remodeling

RNA-seq analysis of FACS-isolated Sun1-positive PT cells identified canonical Wnt/β-catenin signaling as one of the most significantly enriched pathways following *Hnf1b* deletion (Supplemental Figure 2B). Consistent with this enrichment, *Hnf1b*-deficient PT cells exhibited increased expression of multiple Wnt ligand genes, including *Wnt11, Wnt7b, Wnt4, Wnt5a, Wnt5b,* and *Wnt16*, with particularly strong induction of *Wnt11* and *Wnt7b* (Supplemental Table 1). In parallel, mutant kidneys demonstrated increased expression of the canonical Wnt/β-catenin target genes LEF1 and PAX2 (Figures 6A and 6B) (41, 42). In control kidneys, LEF1 expression was largely restricted to medullary stromal cells, with only rare Sun1-negative cortical cells showing detectable signal (Figure 6A). Following *Hnf1b* deletion, LEF1 expression was markedly expanded within the cortex and was detected in both Sun1-positive PT-derived epithelial cells and adjacent Sun1-negative stromal cells. Similarly, whereas PAX2 expression in control kidneys was strongest in the collecting duct and only weakly detected in Sun1-positive PT cells, *Hnf1b*-deficient Sun1-positive PT cells exhibited strong ectopic PAX2 expression (Figure 6B). Together, these findings indicate that loss of *Hnf1b* induces acquisition of a Wnt ligand-expressing epithelial state accompanied by activation of canonical Wnt/β-catenin-dependent transcriptional programs.

**Figure 6.**
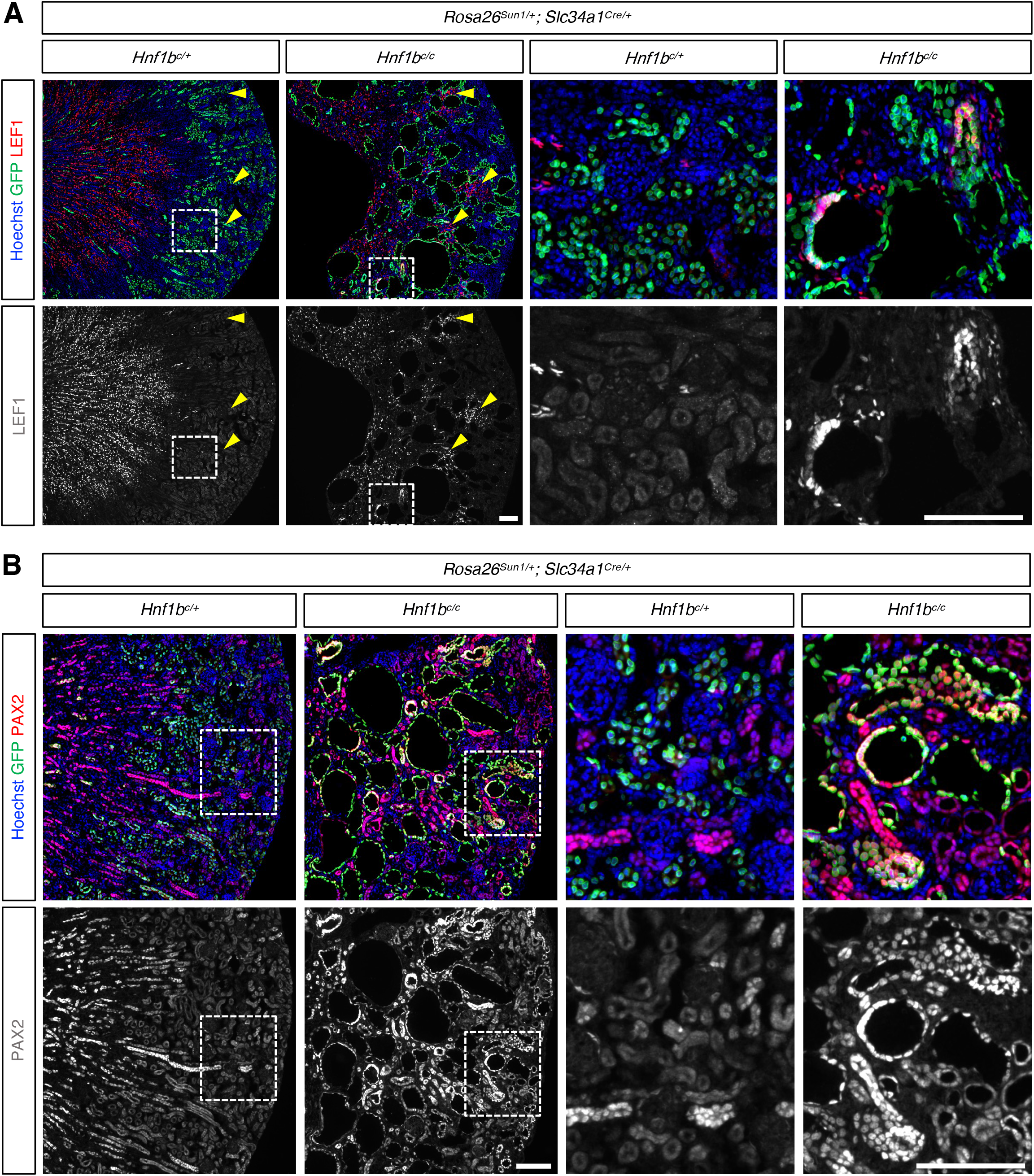
Loss of *Hnf1b* in PT cells activates canonical Wnt target genes in tubular and adjacent stromal cells. (A) In control kidneys, LEF1, a Wnt/β-catenin target gene, is largely restricted to the medulla, whereas mutant kidneys exhibit expanded cortical LEF1 expression in both SUN1+ PT–derived cells and adjacent SUN1− stromal cells. Yellow arrowheads indicate cortical LEF1+ stromal cells. Dotted boxes denote regions shown at higher magnification in the corresponding panels on the right. (B) PAX2, another Wnt/β- catenin target gene, is ectopically expressed in *Hnf1b*-deficient SUN1+ PT cells compared with controls. Dotted boxes denote regions shown at higher magnification in the corresponding right panels. (A, B) Representative images from four biological replicates per genotype are shown. Stage: postnatal day 9; Scale bar: 100 μm.

Activation of Wnt signaling was not restricted to PT-derived epithelial cells but was also associated with remodeling of adjacent stromal populations. While PBX1-positive stromal cells were only sparsely present in the cortex of control kidneys, they were markedly expanded following *Slc34a1Cre*-mediated *Hnf1b* deletion (Figure 7A). Mutant kidneys also exhibited increased expression of PDGFRB in stromal cells together with enhanced interstitial collagen deposition detected by Picrosirius Red staining (Figure 7B and Supplemental Figure 8), indicating stromal expansion and progressive interstitial fibrosis. Collectively, these results demonstrate that loss of *Hnf1b* in PT cells activates Wnt/β-catenin-associated transcriptional programs within both epithelial and stromal compartments and promotes remodeling of the renal interstitium.

**Figure 7.**
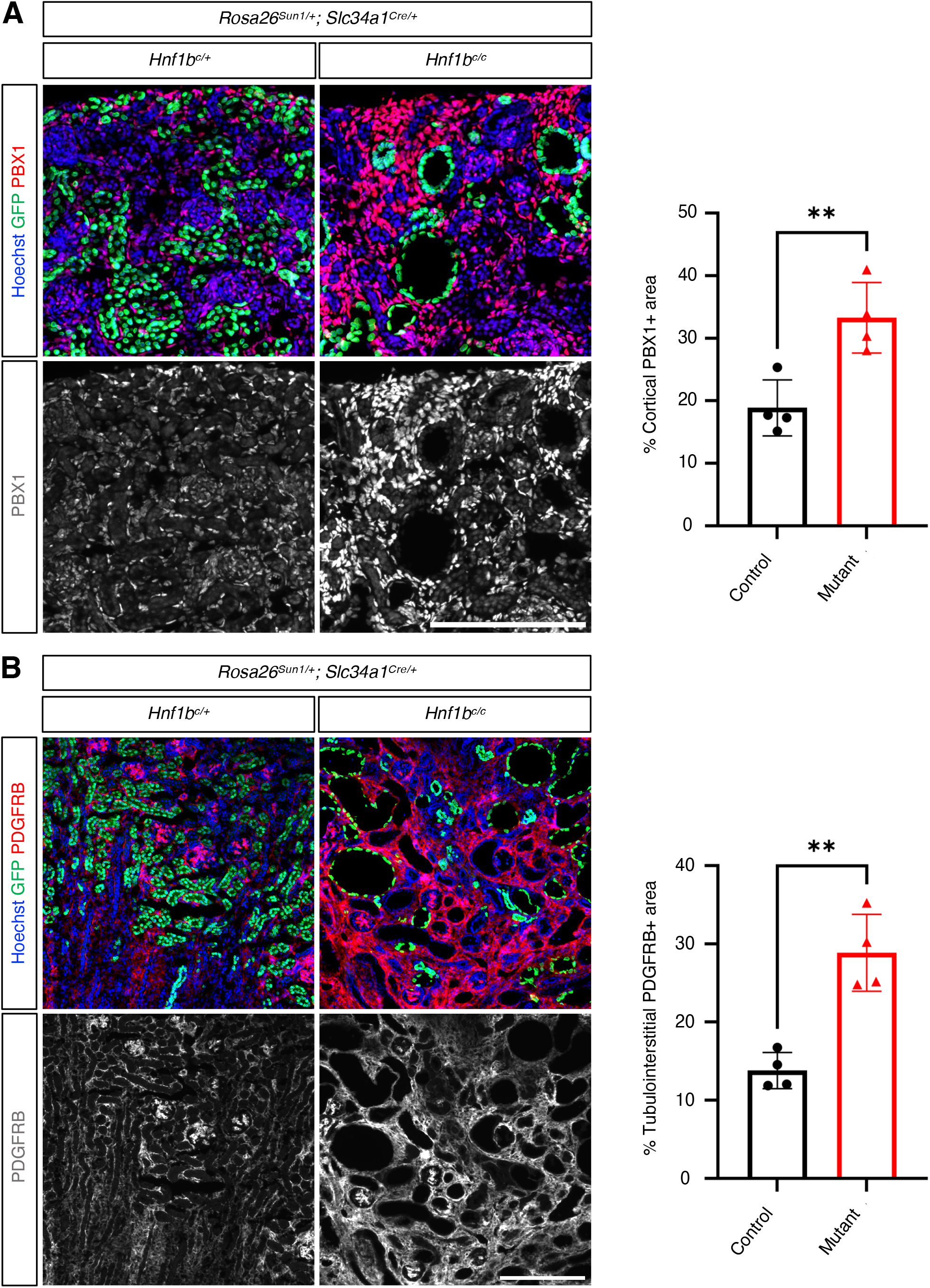
***Hnf1b* deficiency in PT cells promotes expansion of the cortical stromal compartment.** (A) PBX1-positive stromal cells are markedly expanded within the cortical interstitium of *Hnf1b* mutant kidneys and accumulate around renal tubules. In control kidneys, PBX1 expression is largely restricted to the medulla, with only rare PBX1-positive stromal cells present in the cortex. Quantification revealed a significant increase in PBX1-positive cortical area in mutant kidneys compared with controls. (B) In control kidneys, PDGFRB expression is largely confined to sparse interstitial cells, with additional staining in glomeruli and Bowman’s capsule. In contrast, *Hnf1b* mutant kidneys exhibit marked expansion of PDGFRB-positive cells throughout the cortical tubulointerstitium. Quantification demonstrated a significant increase in PDGFRB-positive interstitial area in mutant kidneys compared with controls. (A, B) Four biological replicates were analyzed per genotype. Each dot represents one animal, and the images are representative of these samples. Data are presented as mean ± SEM. Statistical significance was assessed using an unpaired two-tailed Student’s *t* test. **P< 0.01. Stage, postnatal day 9; Scale bars, 100 μm.

## DISCUSSION

Our findings demonstrate that *Hnf1b* is required not only for nephron patterning during development but also for active maintenance of PT identity after segmental identity has been established. Using a post-specification deletion strategy, we show that loss of *Hnf1b* in established PT cells results in marked disruption of PT-specific transcriptional programs and ectopic induction of podocyte-associated genes, revealing a previously unappreciated degree of segmental plasticity within differentiated nephron epithelia. Importantly, activation of podocyte-associated genes in *Hnf1b*-deficient PT cells indicates engagement of an alternative nephron segment program rather than generalized dedifferentiation. Together, these findings establish *Hnf1b* as a critical regulator of PT identity and support a model in which differentiated nephron epithelia require continuous transcriptional enforcement to maintain segment-specific identity and suppress alternative nephron segment programs.

Loss of *Hnf1b* resulted in reduced expression of key PT identity genes, including *Hnf4a,* and widespread suppression of PT transport and metabolic programs. Previous studies showed that developmental *Hnf1b* haploinsufficiency delays PT differentiation and reduces expression of PT-enriched transport and metabolic genes (43), while transcriptional profiling of *Hnf1b*-deficient zebrafish PTs revealed broad suppression of solute-transporter genes (44). More recently, deletion of *Hnf1b* in renal tubular epithelial cells was shown to induce loss of epithelial differentiation and CKD-like pathology in adult kidneys, further supporting a postdevelopmental role for *Hnf1b* in maintenance of tubular epithelial homeostasis (24). Given that *Hnf4a* governs the transcriptional networks that define PT differentiation and function (6, 21, 45), loss of *Hnf4a* expression likely represents a key mechanistic link between *Hnf1b* deficiency and disruption of the PT differentiation program. Importantly, the *Hnf1b–Hnf4a* regulatory axis may also explain defects observed in the medullary DTL. Our previous study demonstrated that a subset of DTL cells is derived from the PT lineage and that *Hnf4a* is required for *Aqp1* expression in the DTL, suggesting that *Hnf4a* acts during a PT-to-DTL transitional stage rather than in mature DTL cells (34). Consistent with this model, PT-derived DTL cells lose AQP1 expression following *Hnf1b* deletion, indicating impaired maintenance of DTL identity. Thus, disruption of *Hnf4a*-dependent transcriptional programs established in PT-derived precursor cells likely contributes to both PT and DTL abnormalities in *Hnf1b*-deficient kidneys.

Our findings also reveal an unanticipated role for *Hnf1b* in suppressing podocyte gene programs within differentiated PT cells. In zebrafish pronephros, knockdown of the *Hnf1b* orthologs *hnf1aa* and *hnf1ab* results in expansion of the *wt1* expression domain into the region normally occupied by the PT segment, leading to the conclusion that *Hnf1b* suppresses podocyte fate during nephrogenesis (13). Because conditional deletion of *Hnf1b* in mice using *Six2Cre* or *Wnt4Cre* did not induce ectopic *Wt1* expression (14, 16, 17), the zebrafish phenotype was generally interpreted as a developmental consequence of defective nephron patterning. Our results challenge this interpretation. By deleting *Hnf1b* after PT segmental identity is established, we demonstrate that differentiated mouse PT cells activate podocyte-associated genes, including *Wt1* and *Nphs1*, following loss of *Hnf1b*. These findings suggest that the zebrafish and mouse phenotypes reflect a conserved function of *Hnf1b* in suppressing podocyte transcriptional programs within PT cells. Importantly, because early deletion models disrupt PT formation itself, they could not reveal this post-specification role.

In addition to transcriptional reprogramming, *Hnf1b* deficiency disrupted epithelial organization. Loss of *Hnf1b* resulted in reduced expression of E-cadherin (CDH1), induction of the mesenchymal markers VIM and TAGLN, and defects in apical-basal polarity. *HNF1B* loss has been shown to activate mesenchymal and profibrotic programs in renal epithelial cells (25). In ureteric bud–derived epithelia, *Hnf1b* deletion also disrupted cell–cell contacts, apicobasal polarity, and basement membrane organization (46). Similar abnormalities have been described in cyst-lining epithelia in polycystic kidney disease, suggesting that epithelial destabilization is a common feature of cystogenesis (38, 47, 48). Despite these abnormalities, *Hnf1b*-deficient cells remained organized as epithelial structures lining tubules and cysts, indicating partial destabilization of epithelial identity rather than complete epithelial collapse. These observations suggest that *Hnf1b* coordinates both the transcriptional and structural components of PT differentiation and that loss of *Hnf1b* creates a cellular environment permissive for cyst initiation.

In addition to disrupting PT identity and epithelial organization, *Hnf1b* deficiency was associated with increased epithelial proliferation and activation of developmental growth and signaling programs. Mutant kidneys exhibited increased Ki67 labeling together with induction of *Igf2*, a fetal growth factor that is highly expressed during nephrogenesis and subsequently downregulated following kidney maturation (49–51). Loss of *Hnf1b* was also accompanied by increased expression of multiple Wnt ligands, particularly *Wnt7b* and *Wnt11*, together with elevated expression of canonical Wnt/β-catenin target genes including *Lef1* and *Pax2*. The induction of Lef1 is consistent with previous evidence that HNF1B restrains canonical Wnt signaling through direct transcriptional repression of *Lef1* in renal epithelial cells (52). Because both IGF2 and Wnt signaling regulate cellular growth during kidney development, reactivation of these pathways may contribute to the hyperproliferative phenotype observed in *Hnf1b*-deficient PT cells. Activation of Wnt signaling was not restricted to the epithelium. Expansion of LEF1+ and PBX1+ stromal populations, together with increased numbers of PDGFRB-positive cells and enhanced interstitial collagen deposition, indicates activation of developmental epithelial-stromal signaling pathways and progressive interstitial remodeling. Our findings are consistent with recent evidence that HNF1B loss in adult renal tubular cells causes interstitial fibrosis and progressive CKD (24). These observations are also notable in light of the tubulointerstitial changes observed in patients with RCAD, in whom heterozygous HNF1B mutations are associated with hyperuricemic nephropathy and autosomal dominant tubulointerstitial kidney disease (53). Together, these findings suggest that the interstitial remodeling observed in our model may reflect a conserved pathological response to HNF1B dysfunction. Similar epithelial-stromal Wnt interactions have been implicated in renal fibrosis, where epithelial-derived Wnt ligands activate β-catenin signaling in neighboring stromal cells (54). Together with previous studies demonstrating that pharmacologic or genetic inhibition of canonical Wnt/β-catenin signaling ameliorates cystogenesis in mouse models of polycystic kidney disease (55), our findings support a model in which loss of *Hnf1b* reactivates developmental growth and signaling pathways, including *Igf2* and Wnt/β- catenin signaling, that contribute to epithelial proliferation, stromal remodeling, and cyst progression.

Overall, our findings demonstrate that mature nephron segments require continuous activity of identity-maintaining transcription factors to preserve their differentiated state and suppress alternative segment-specific programs. By extending the role of *Hnf1b* beyond early nephron patterning, this work identifies active maintenance of nephron segment identity as a fundamental requirement for epithelial homeostasis and reveals how disruption of this process can lead to epithelial destabilization, developmental pathway reactivation, and cystogenesis.

## METHODS

### Sex as a biological variable

Both male and female mice were used in this study, with equal representation of each sex. Mutant and control animals of both sexes were included in all analyses unless otherwise indicated. Sex-based differences were assessed, and no appreciable differences were observed between male and female mice across the measured outcomes. Therefore, data from male and female animals were pooled for subsequent analyses.

### Generation of the conditional *Hnf1b* (*Hnf1b^c^*) mouse line

A conditional *Hnf1b* allele was generated by flanking exon 2 with loxP sites using CRISPR/Cas9–mediated genome editing, resulting a frameshift and loss of Hnf1b function upon Cre recombination. Two single-guide RNAs targeting intronic regions upstream (Sg299; ACTGTTGCTTGTGTCCCAGG) and downstream (Sg300; GTGTAACTACTTGTGTAGCA) of exon 2 were used together with single-stranded donor oligonucleotides containing loxP sequences. CRISPR reagents were delivered to fertilized zygotes by sequential electroporation, and edited embryos were transferred into pseudopregnant CD-1 females. Offspring were screened for loxP site insertion by PCR and validated by Sanger sequencing. One founder line (#1665) carrying both loxP sites was used to establish the *Hnf1b* conditional (*Hnf1b^c^*) mouse line. Genotyping PCR was performed using CCTTAGTTTCCCCTCCACCAAGA and CACCCTGAAGGCCTTTTCAGTGT. This reaction yields a 142-bp amplicon for the wild-type *Hnf1b* allele and a 200-bp amplicon for the conditional *Hnf1b* allele.

#### Mice and breeding strategy

PT-specific deletion of *Hnf1b* was achieved using the *Slc34a1eGFPCre* (*Slc34a1Cre*; JAX: 040319) allele, which drives constitutive Cre expression in differentiated PT cells (34). For lineage tracing and isolation of recombined PT cells, we used the *Rosa26-Sun1* (JAX: 021039) and *Rosa26-NuTRAP* (JAX: 029899) reporter alleles. *Rosa26-Sun1* expresses a SUN1-superfolder GFP fusion protein at the nuclear envelope following Cre-mediated recombination (56), enabling lineage tracing and histologic analysis of recombined cells. *Rosa26-NuTRAP* expresses EGFP-tagged ribosomal protein L10a following Cre-mediated recombination (57) and was used for fluorescence-activated cell sorting (FACS) and bulk RNA-seq analysis. For lineage-tracing experiments, *Slc34a1^Cre/Cre^; Hnf1b^c/+^* mice were crossed with *Hnf1b*^c/c^; *Rosa26^Sun1/Sun1^*mice to generate *Hnf1b* mutant mice (*Slc34a1^Cre/+^; Hnf1b*^c/c^; *Rosa26^Sun1/+^*) and littermate controls (*Slc34a1^Cre/+^*; *Hnf1b^c/+^*; *Rosa26^Sun1/+^*). For FACS and bulk RNA-seq experiments, the *Rosa26-Sun1* allele was replaced with *Rosa26-NuTRAP*. All mice in this report were bred and maintained in a mixed genetic background. Animals were centrally managed by Northwestern University Center for Comparative Medicine (CCM), an animal care facility that operates within full compliance of the NIH guidelines. Mice were monitored daily and housed in a controlled environment with a 12-hour light/12-hour dark cycle, with ad libitum access to water and a standard chow diet. At the first sign of pain, suffering or distress, mice were euthanized. Humane end points were established to include lethargy, labored breathing, or swelling.

#### Fluorescence-activated cell sorting (FACS) and bulk RNA-seq

FACS and bulk RNA sequencing were performed on *Hnf1b* mutant and control kidneys, wherein *Slc34a1Cre* was used to activate the *Rosa26-NuTrap* reporter and delete the conditional *Hnf1b* allele. Kidneys at postnatal day 1 (P1) were dissociated using TrypLE Select Enzyme 10X (Gibco) and mechanical pipetting. Cells were washed, resuspended in PBS containing 1% FBS and 10 mM EDTA, and filtered through 40 μm nylon strainers. GFP-positive cells were isolated using a BD FACSAria SORP system at the Northwestern University RHLCCC Flow Cytometry Facility, supported by NIH grants 1S10OD011996-01 and 1S10OD026814-01. mRNA was isolated from total RNA using the NEBNext Poly(A) mRNA Magnetic Isolation Module (E7490L, New England Biolabs). Fragmentation of mRNA and cDNA synthesis were performed using NEBNext RNA First Strand Synthesis Module (E7525L) and NEBNext RNA Second Strand Synthesis Module (E6111L). The cDNA’s were processed to sequencing libraries using ThruPLEX DNA-seq 12S Kit (R400675 and R400695, Takara). Libraries were sequenced on Illumina NovaSeq X Plus or Element AVITI at the NUSeq Core facility at Northwestern University.

#### RNA-seq analysis

Paired-end RNA-seq reads were mapped to the UCSC mouse reference genome (mm10) using the STAR aligner (58). Only uniquely aligned reads were used for differential gene analysis. Gene-level read counts were obtained using FeatureCounts, with the following parameters: “-s 2 -O --fracOverlap 0.8” (59). Differential expression analysis was conducted using the DESeq2 package (60). Genes exhibiting an absolute fold-change > 2.0 and FDR < 0.01 were considered significantly differentially expressed. Gene ontology (GO) analysis was performed using EnrichR (61). Gene expression profiles of selected genes were visualized using single-cell RNA-seq datasets from mouse kidneys at embryonic day 18.5 and postnatal day 0 (GSE214024 and GSE275601) (34, 35). Dot plots and module score projections were generated in Seurat (62). Module scores for bulk RNA-seq differentially expressed genes were calculated using the AddModuleScore function and projected onto the UMAP embedding.

#### Immunofluorescence staining and microscopy

Kidneys from mutant and control animals of both sexes were analyzed. For immunofluorescence analyses, biological replicates consisted of kidneys collected from independent animals; two control and four mutant biological replicates were examined, unless otherwise indicated. Tissue preparation, staining and imaging were performed in parallel for each experiment to minimize technical variation. Sample size was determined by litter size and breeding outcomes. No formal sample size calculation was performed. No exclusion criteria were defined, and no animals or data points were removed from the analysis. Samples were processed in randomized order, and control and mutant kidneys were embedded within the same block to reduce batch effects. The investigator was aware of sample genotypes during processing and analysis. Kidneys were fixed in phosphate-buffered saline (PBS) containing 4% paraformaldehyde (PFA) for 20 minutes and then incubated overnight at 4°C in PBS containing 10% sucrose. Tissues were embedded in OCT compound (Thermo Fisher Scientific) and stored at -80°C. Cryosections (8 μm) were incubated overnight at 4°C in PBS containing 0.1% Triton X-100, 5% heat-inactivated sheep serum, and primary antibodies (Supplemental Table 2). Fluorophore-labeled secondary antibodies were sourced from Invitrogen or Jackson ImmunoResearch (Supplemental Table 2). All images were obtained using a Nikon Ti2 widefield microscope.

#### Hybridization Chain Reaction

Hybridization chain reaction (HCR) was performed as previously described with minor modifications (63, 64). Cryosections were washed in 1× PBS and subjected to autofluorescence bleaching using a combined light and hydrogen peroxide treatment. Sections were illuminated with a gooseneck LED desk lamp (5 V, 2 A output) in the presence of an H_₂_O_₂_-based chemical bleaching solution (C762T82, H1070-100ML-CS6; Spectrum Chemicals; Supplemental Table 3) at 4°C for 1 hour. After bleaching, sections were washed at room temperature in 1× PBS (3 × 10 minutes), PBS containing Tween-20 (3 × 5 minutes), and 5× saline sodium citrate with Tween-20 (5× SSC-T; 3 × 5 minutes), followed by a brief rinse in 1× PBS. Sections were then incubated in probe hybridization buffer for 30 minutes at 37°C, followed by overnight hybridization at 37°C with ssDNA probe sets targeting the following mouse transcripts: *Mafb* (NM_010658), *Cldn5* (NM_013805), *Nphs1* (NM_019459), *Plat* (NM_008872), and *Igf2* (NM_001122737.2). The following day, sections were incubated in probe wash buffer for 30 minutes at 37°C, followed by two additional washes in probe wash buffer (2 × 15 minutes at 37°C) and washes in 5× SSC-T (3 × 5 minutes at room temperature). For signal amplification, sections were equilibrated in amplification buffer for 30 minutes at room temperature. The buffer was then replaced with amplification buffer containing Amplifier B1 DNA hairpins diluted according to the manufacturer’s instructions, and sections were incubated in the dark to allow hybridization chain reaction–mediated signal amplification. Slides were subsequently washed in 1× PBS (5 minutes) and 5× SSC-T (1 × 5 minutes followed by 2 × 30 minutes) in the dark. Nuclei were counterstained with Hoechst (1:2000 dilution; Invitrogen) for 10 minutes. After mounting, slides were kept at room temperature and protected from light until imaging. Detailed information regarding target genes, probe sequences, and amplifier hairpins is provided in Supplemental Table 4.

#### Histology and Picrosirius Red staining

Kidneys were fixed in 4% paraformaldehyde (PFA) in PBS overnight and submitted to the Mouse Histology & Phenotyping Laboratory (MHPL) at the Robert H. Lurie Comprehensive Cancer Center of Northwestern University, which is supported by the National Cancer Institute (NCI) P30CA060553. Paraffin sections (3 μm) were stained with hematoxylin and eosin or Picrosirius Red. Brightfield images were acquired using a Nikon Ti2 widefield microscope housed in the Center for Advanced Microscopy/Nikon Imaging Center (RRID: SCR_020996) at Northwestern University. For quantification of collagen deposition, Picrosirius Red–positive area was measured in ImageJ/Fiji using a uniform threshold applied to all images within the same experiment. Collagen deposition was expressed as the percentage of Picrosirius Red–positive area relative to the total analyzed tissue area. Measurements from multiple sections were averaged to generate one value per animal. Quantification was performed blinded to genotype.

#### Image and cell quantification

PDGFRB-positive, PBX1-positive, and *Igf2*-positive areas in kidney sections were quantified using ImageJ/Fiji from images acquired under identical imaging settings (65). For each marker, a single threshold was applied uniformly to all images within the same experiment, including control and mutant samples, to define positive signal. Positive area was quantified using the ImageJ area fraction function and expressed as the percentage of the analyzed region of interest (ROI). *Igf2* signal was quantified from HCR images, whereas PDGFRB and PBX1 signals were quantified from immunostained kidney sections. Glomeruli, large vessels, and tissue-processing artifacts were excluded from analysis. PDGFRB-positive area was quantified within the cortical tubulointerstitium, whereas PBX1-positive area and *Igf2* signal were quantified within the renal cortex. Approximately three nonoverlapping cortical fields measuring 389.6 × 389.6 μm were analyzed per animal across multiple kidney sections. Field-level measurements were averaged to generate one value per animal. Four animals were analyzed per genotype, with each animal treated as one biological replicate. Quantification was performed in a blinded manner. Cell proliferation was assessed by immunofluorescent staining for Ki67. Approximately three nonoverlapping fields measuring 389.6 × 389.6 μm were analyzed per animal across multiple kidney sections using QuPath (v0.5.1) (66). The numbers of GFP-positive cells and GFP/Ki67 double-positive cells were determined for each field, and the proliferation index was calculated as the percentage of GFP/Ki67 double-positive cells relative to the total number of GFP-positive cells. Quantification was performed in a blinded manner.

#### Statistical analyses

All statistical analyses were performed using GraphPad Prism. Comparisons between two groups were conducted using unpaired two-tailed Student’s *t*-tests. Sample sizes were not predetermined using statistical methods, and data distributions were not formally assessed for normality. Data are presented as mean ± SEM. Statistical significance was defined as *P* < 0.05 (*), *P* < 0.01 (**), and *P* < 0.001 (**\*\*\***).

#### Study approval

All animal experiments were approved by the Institutional Animal Care and Use Committees (IACUC) at Northwestern University and Cincinnati Children’s Hospital Medical Center and were conducted in accordance with the NIH Guide for the Care and Use of Laboratory Animals.

## Data availability

The bulk RNA sequencing data generated in this study have been deposited in the Gene Expression Omnibus (GEO) under accession number GSE318534. All other data supporting the findings of this study are available from the corresponding author upon reasonable request.

## Supporting information

RNA-seq analysis of Hnf1b by Slc34a1Cre mutant and control kidneys at P1

Primary and secondary antibodies

HCR RNA FISH solutions

HCR probs and amplifiers

## Author Contributions

ZDG and JSP designed the study. ZDG, EC, and JSP conducted experiments. MS, CA, and HWL performed bioinformatic data analysis. ZDG and JSP curated and validated the data and prepared the figures. YCH and AH contributed to methodology development. JSP acquired funding, provided resources, and administered the project. ZDG and JSP wrote the original draft of the manuscript, and ZDG, EC, and JSP reviewed and edited the final version. All authors approved the final version of the manuscript.

## Acknowledgments

We thank the Robert H. Lurie Comprehensive Cancer Center of Northwestern University in Chicago, IL, for the use of the Flow Cytometry Core Facility, which provided cell sorting services. The Lurie Cancer Center is supported in part by NCI Cancer Center Support Grant P30 CA060553. We also thank the Northwestern University NUSeq Core Facility for sequencing services.

## Supplemental Material

Supplemental Figure 1. Integrating Bulk RNA-seq differential expression with single-nucleus RNA-seq UMAP to identify cell-type-specific responses to Hnf1b deletion in PT cells.

Supplemental Figure 2. Gene Ontology and KEGG pathway enrichment analyses of differentially expressed genes in Hnf1b mutant kidneys.

Supplemental Figure 3. Loss of *Hnf1b* in PT cells leads to ectopic expression of podocyte-specific markers.

Supplemental Figure 4. S*l*c34a1Cre -mediated *Hnf1b* deletion causes early medullary cyst formation

Supplemental Figure 5. Loss of *Hnf1b* disrupts AQP1 expression in PT-derived descending thin limb cells lining medullary cysts.

Supplemental Figure 6. Loss of *Hnf1b* impairs apical–basal polarity in PT cells.

Supplemental Figure 7. Loss of *Hnf1b* in PTs increases epithelial proliferation and induces *Igf2* expression.

Supplemental Figure 8. Loss of *Hnf1b* in PT cells increases interstitial collagen deposition.

## Supplemental Methods

Supplemental Table 1. RNA-seq analysis of *Hnf1b* by *Slc34a1Cre* mutant and control kidneys at P1

Supplemental Table 2. Primary and secondary antibodies used in this study

Supplemental Table 3. HCR RNA FISH solutions Supplemental Table 4. HCR probs and amplifiers

