## Supplemental Figures for "Loss of *Hnf1b* in differentiated proximal tubule cells uncovers nephron segment plasticity"

### Supplemental Figure 1

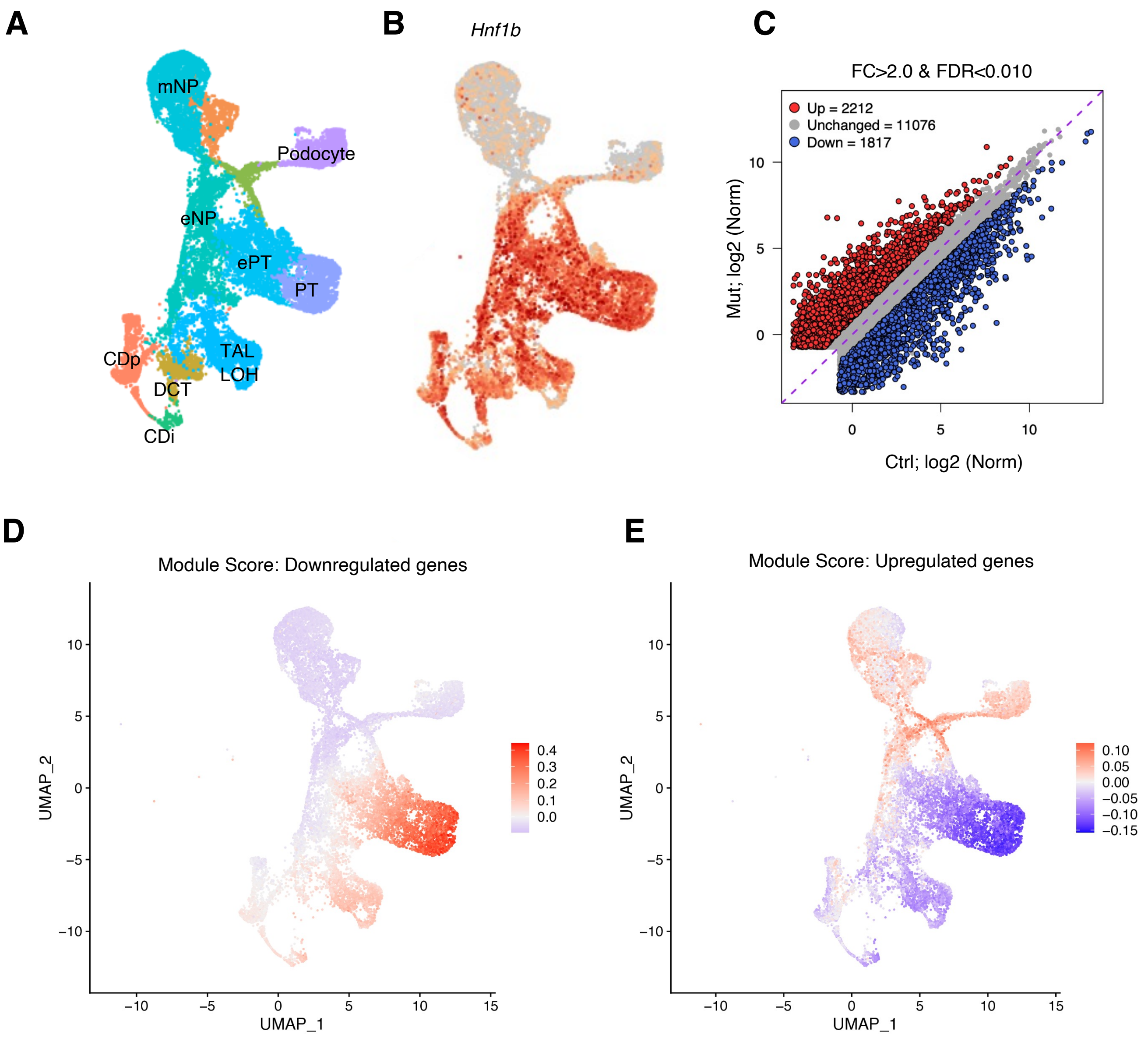


**Supplemental Figure 1. Differentially expressed genes in *Hnf1b*-deficient proximal tubule cells map predominantly to proximal tubule and podocyte populations in a wild-type single-cell RNA-seq atlas.** (A) UMAP visualization of single-cell RNA-seq data from mouse kidneys at E18.5 and P0 (GSE214024 and GSE275601) after subsetting to nephron and collecting duct lineages. mNPs, mesenchymal nephron progenitors; eNPs, epithelial nephron progenitors; ePT, early proximal tubules; PT, proximal tubules; TAL LOH, thick ascending limb of the loop of Henle; DCT, distal convoluted tubule; CDi, collecting duct intercalated cells; CDp, collecting duct principal cells. (B) Feature plot of *Hnf1b* expression showing broad expression throughout nephron and collecting duct lineages, with exclusion from mNPs and podocytes. (C) Scatter plot showing differentially expressed genes (DEGs) identified by bulk RNA-seq of FACS-isolated PT cells from *Hnf1b* mutant and control kidneys. Red and blue dots indicate genes that are upregulated and downregulated, respectively, in *Hnf1b*-deficient PT cells. Gray dots represent genes that are not differentially expressed. (D) UMAP projection of module scores for genes downregulated in *Hnf1b*-deficient PT cells. Color intensity reflects module score, with enrichment predominantly localized to the PT cluster, consistent with a requirement for *Hnf1b* in maintaining PT identity. (E) UMAP projection of module scores for genes upregulated in *Hnf1b*-deficient PT cells. Color intensity reflects module score, with enrichment predominantly localized to the podocyte cluster, indicating ectopic activation of podocyte-associated gene programs following PT-specific *Hnf1b* deletion

### Supplemental Figure 2

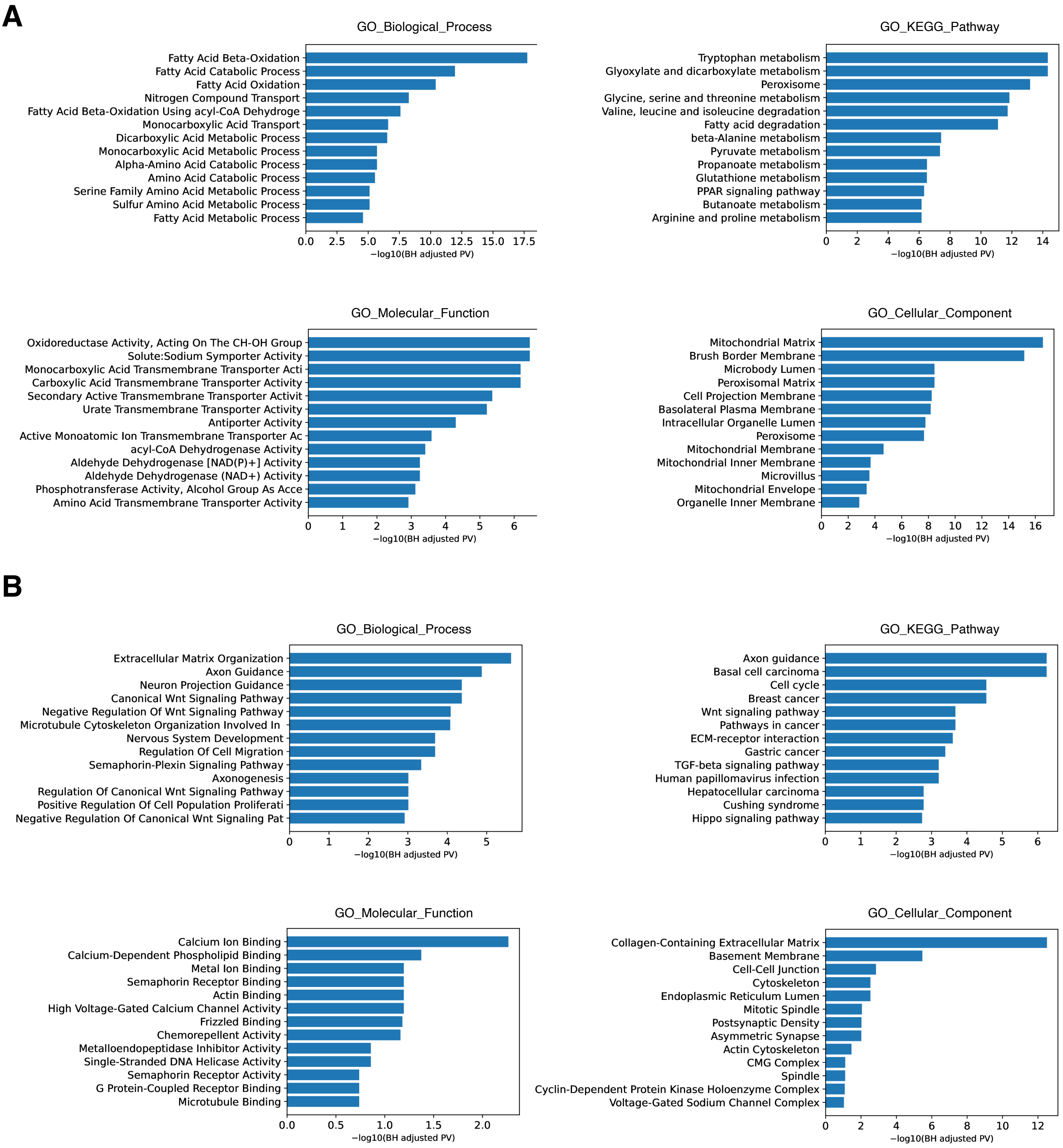


**Supplemental Figure 2. Gene Ontology and KEGG pathway enrichment analyses of differentially expressed genes in *Hnf1b*-deficient PT cells.** Bar plots show the top significantly enriched terms (ranked by $-\log_{10}$ adjusted $P$ value), from Gene Ontology (GO; Biological Process, Molecular Function, and Cellular Component) and KEGG pathway analyses (adjusted $P < 0.01$). (A) Downregulated genes ($n$ = 1,817) are enriched for metabolic and solute transport processes characteristic of PT function. (B) Upregulated genes ($n$ = 2,212) are enriched for processes related to extracellular matrix organization, axon guidance, canonical Wnt signaling, and neuronal system development, indicating activation of developmental and cell-cell communication programs.

### Supplemental Figure 3

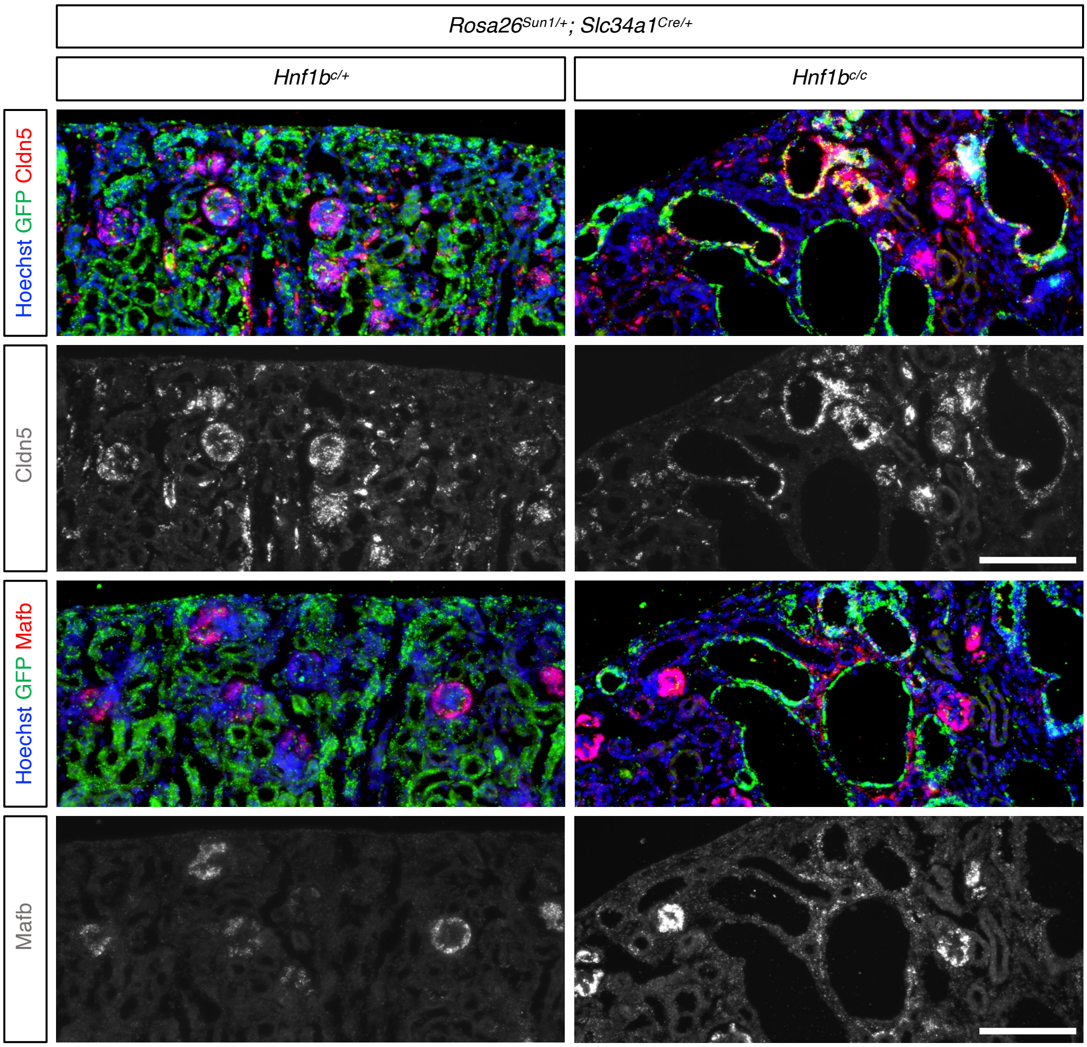


**Supplemental Figure 3. Loss of *Hnf1b* in proximal tubule cells leads to ectopic expression of podocyte-associated markers.** HCR-FISH demonstrates ectopic expression of *Cldn5* and *Mafb* in GFP-positive *Hnf1b*-deficient PT cells. *Cldn5* encodes the tight junction protein Claudin-5, which is primarily expressed in podocytes and endothelial cells, whereas *Mafb* encodes a transcription factor selectively expressed in podocytes. In control kidneys, GFP-positive PT cells show little or no expression of *Cldn5* or *Mafb*. In contrast, *Hnf1b* mutant kidneys exhibit expression of both genes in podocytes and a subset of GFP-positive PT cells. Representative images from four biological replicates per genotype are shown. Stage: postnatal day 9; Scale bar: 100 µm.

### Supplemental Figure 4

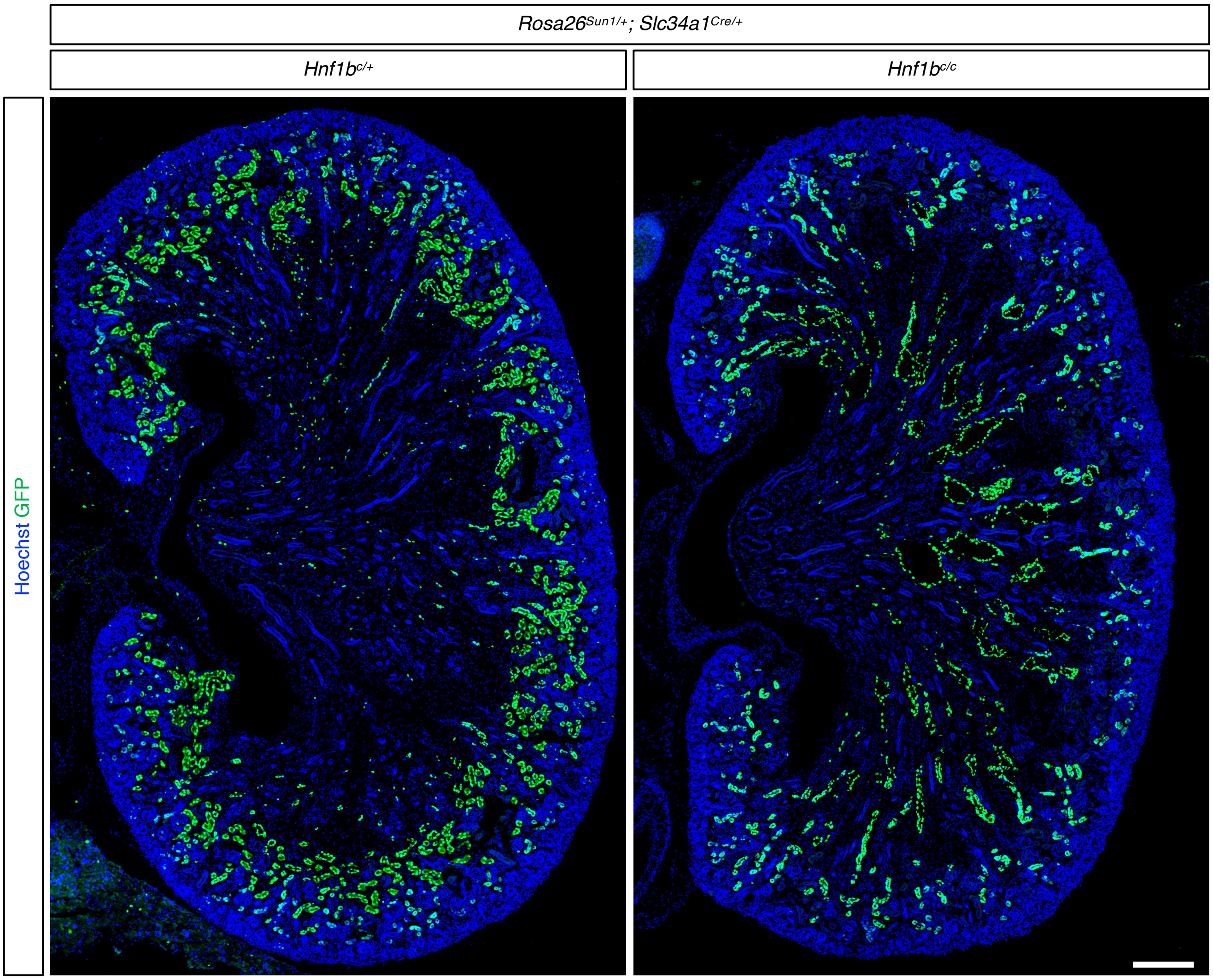


**Supplemental Figure 4. *Slc34a1Cre*-mediated *Hnf1b* deletion causes early medullary cyst formation.** *Slc34a1Cre* labels all PT cells and a subset of descending limb cells with the Rosa26-Sun1 (GFP) reporter. At postnatal day 1, most GFP-positive cells in the cortex of mutant kidneys are non-cystic, whereas GFP-positive cells in the medulla exhibit prominent cystic dilations. These findings indicate that cystogenesis is initiated within PT-derived medullary DTL segments before the appearance of cortical cysts. Representative images from four biological replicates per genotype are shown. Stage: postnatal day 1; Scale bar: 200 µm.

### Supplemental Figure 5

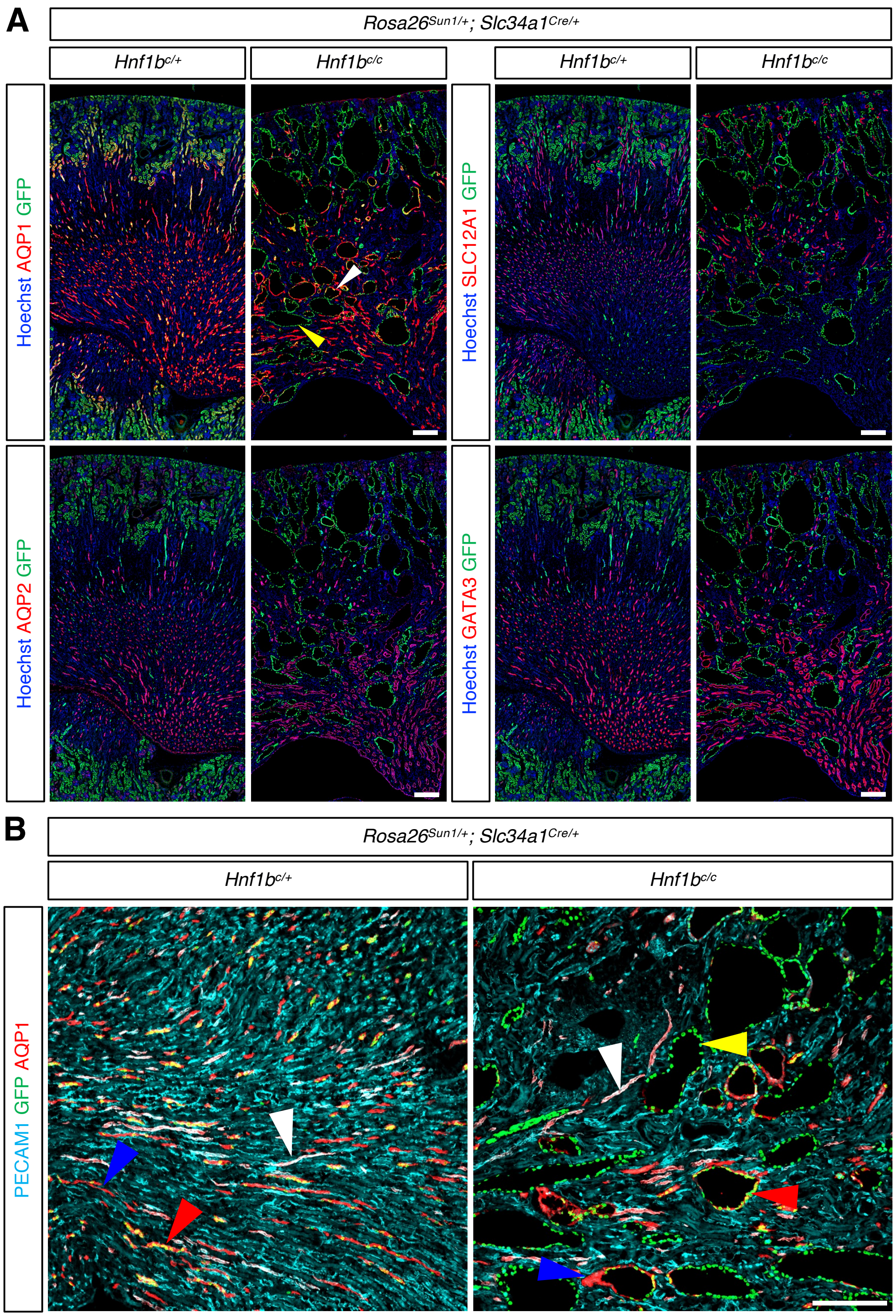


**Supplemental Figure 5. Loss of *Hnf1b* disrupts AQP1 expression in PT-derived descending thin limb (DTL) cells lining medullary cysts.** (A) Medullary cysts in *Hnf1b* mutant kidneys are lined by GFP-positive epithelial cells. A subset of GFP-positive cyst-lining cells retained AQP1 expression (white arrowhead), whereas others lacked detectable AQP1 (yellow arrowheads). Cyst-lining cells were negative for SLC12A1, AQP2, and GATA3, excluding thick ascending limb and collecting duct identities. (B) High-magnification images of the renal medulla stained for AQP1, PECAM1, and GFP. AQP1+/PECAM1+ cells correspond to vasa recta endothelial cells (white arrowheads), whereas AQP1+/PECAM1– cells correspond to epithelial cells of the descending thin limb (DTL). Consistent with our previous lineage-tracing studies, *Slc34a1Cre* mosaically targets PT-derived DTL cells, resulting in both recombined (GFP+) and non-recombined (GFP–) DTL cells in control and mutant kidneys. In control kidneys, all GFP+ DTL cells express AQP1 (red arrowheads). In contrast, *Hnf1b* mutant kidneys contain two populations of GFP+ DTL cells: AQP1+/GFP+ cells that retain AQP1 expression (red arrowheads) and AQP1–/GFP+ cells that have lost detectable AQP1 expression following *Hnf1b* deletion (yellow arrowheads). Non-recombined AQP1+/GFP– DTL cells are indicated by blue arrowheads. These findings indicate that Hnf1b is required for maintenance of AQP1 expression in PT-derived DTL cells. (A, B) Representative images from four biological replicates per genotype are shown. Stage: postnatal day 9; Scale bar: 200 µm.

### Supplemental Figure 6

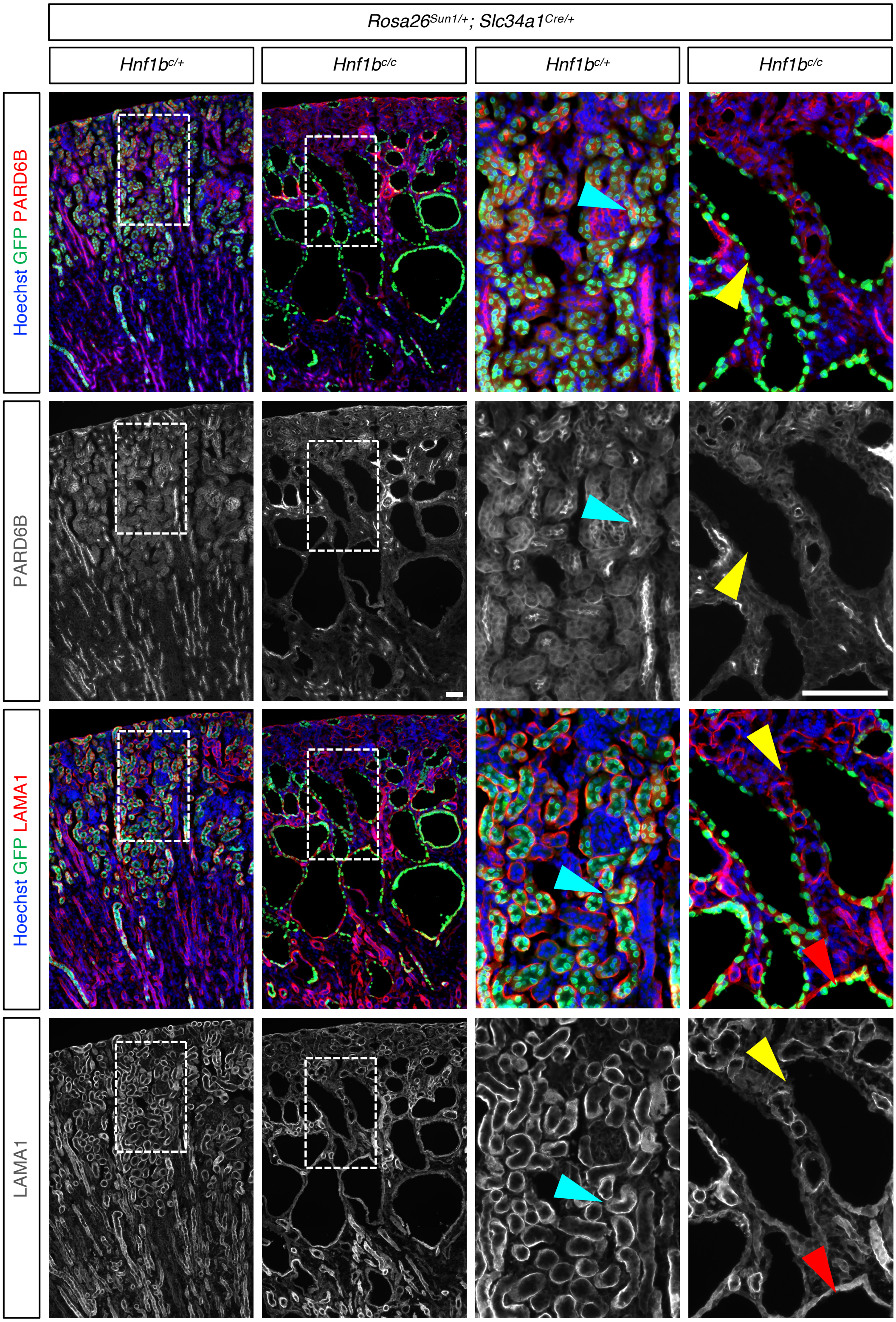


**Supplemental Figure 6. Loss of *Hnf1b* impairs apical–basal polarity in PT cells.** In control kidneys, GFP+ cells display clear apical and basal polarity, marked by PARD6B and LAMA1, respectively. Low-magnification images (left) show overall kidney architecture, and the regions enclosed by white dashed boxes are shown at higher magnification in the corresponding panels on the right. In the high-magnification images, control GFP+ cells exhibit normal apical PARD6B localization and a continuous LAMA1-positive basement membrane (cyan arrowheads). In contrast, GFP+ cells in mutant kidneys lack apical PARD6B expression (yellow arrowheads), indicating disruption of apical polarity. LAMA1 expression in GFP+ mutant cells is either absent (yellow arrowheads) or mislocalized (red arrowheads), consistent with basement membrane disorganization. Representative images from four biological replicates per genotype are shown. Stage: postnatal day 9; Scale bar: 100 µm.

### Supplemental Figure 7

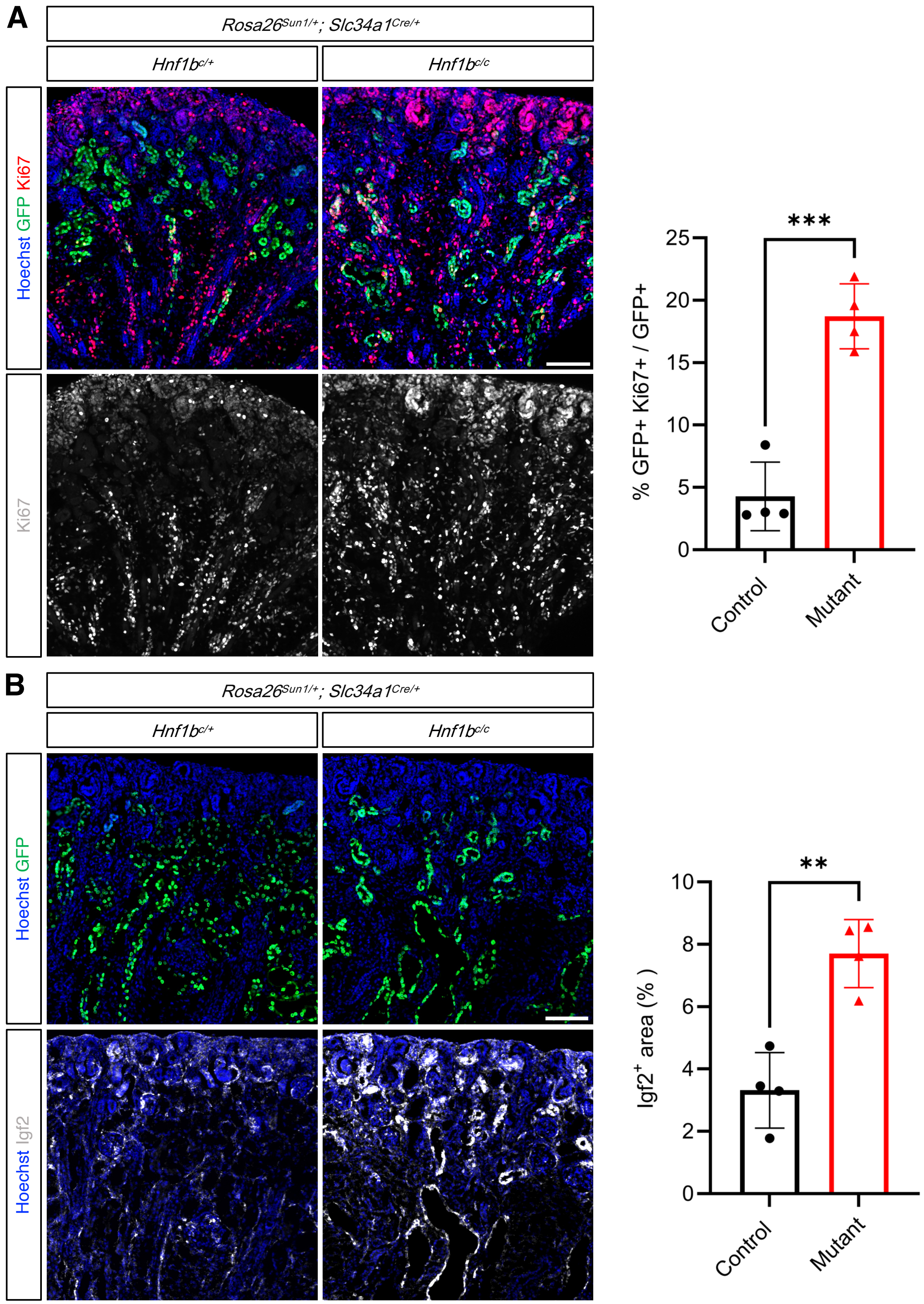


**Supplemental Figure 7. Loss of *Hnf1b* in PTs increases epithelial proliferation and induces *Igf2* expression.** (A) In control kidneys, Ki67-positive cells are sparse within GFP+ PTs. In contrast, *Hnf1b* mutant kidneys exhibit increased Ki67 staining in GFP+ PT cells. Quantification showed a significant increase in the percentage of GFP–positive cells co-expressing Ki67 in mutant kidneys (*** $P < 0.001$). (B) HCR analysis revealed minimal *Igf2* expression in control kidneys, whereas *Hnf1b* mutant kidneys exhibited increased *Igf2* expression. Quantification showed a significant increase in *Igf2* expression in mutant kidneys compared with controls (** $P < 0.01$). (A, B) Four biological replicates were analyzed per genotype. Each dot represents one animal, and the images are representative of these samples. Data are presented as mean ± SEM. Statistical significance was assessed using an unpaired two-tailed Student's *t* test. Stage: postnatal day 1; Scale bar: 100 µm.

### Supplemental Figure 8

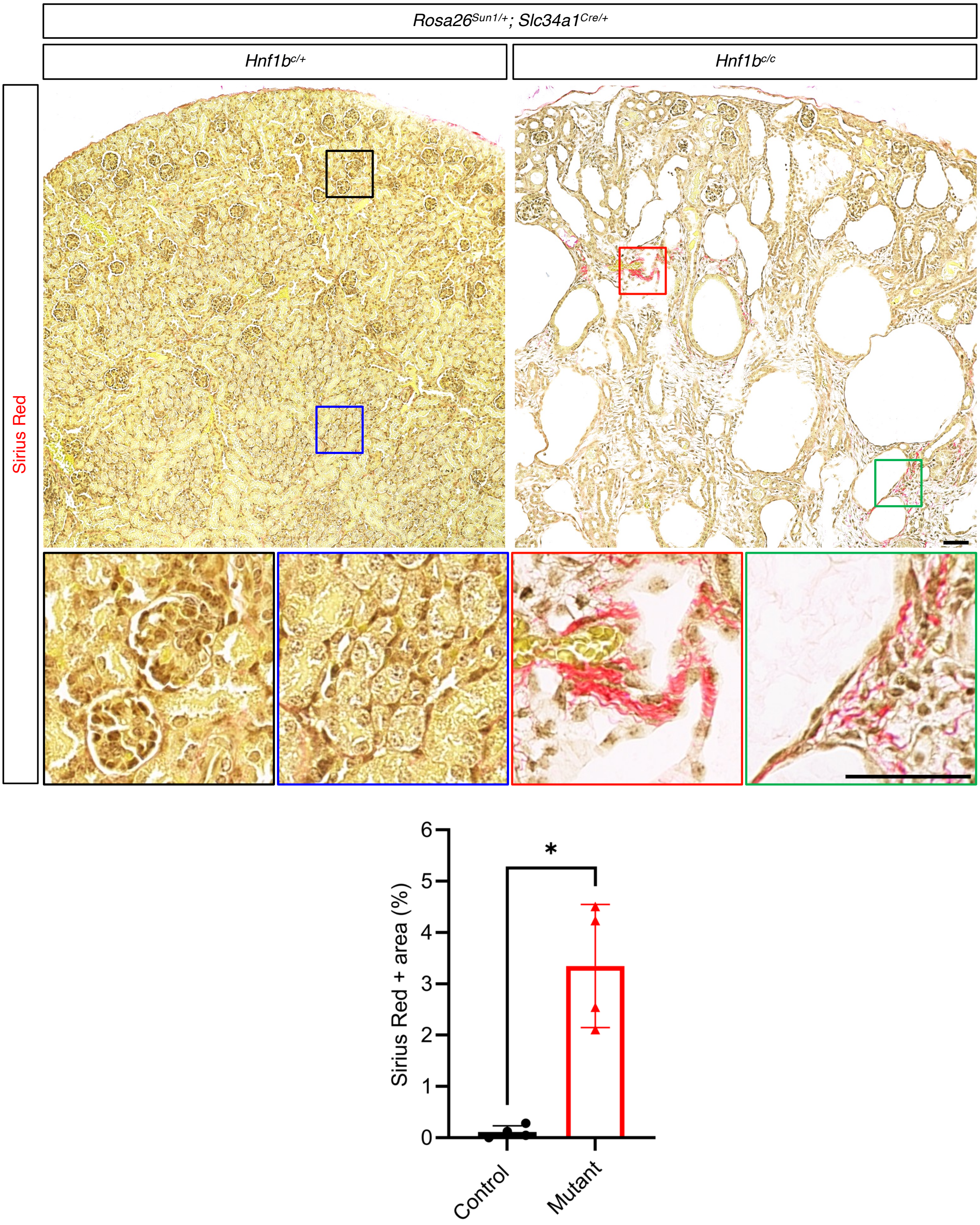


**Supplemental Figure 8. Loss of *Hnf1b* in proximal tubules increases interstitial collagen deposition.** Control kidneys show minimal Picrosirius Red staining, whereas mutant kidneys exhibit increased interstitial collagen deposition, including accumulation around cystic tubules. Insets show higher-magnification views of the boxed regions. Quantification revealed a significant increase in Picrosirius Red–positive area in mutant kidneys compared with controls. Four biological replicates were analyzed per genotype. Each dot represents one animal, and the images are representative of these samples. Data are presented as mean ± SEM. Statistical significance was assessed using an unpaired two-tailed Student's *t* test. $^{*}P < 0.05$. Stage: P9. Scale bars: 100 µm.
